# Predicting Molecular Fingerprint from Electron−Ionization Mass Spectrum with Deep Neural Networks

**DOI:** 10.1101/2020.03.30.017137

**Authors:** Hongchao Ji, Hongmei Lu, Zhimin Zhang

## Abstract

Electron−ionization mass spectrometry (EI-MS) hyphenated gas chromatography (GC) is the workhorse to analyze volatile compounds in complex samples. The spectral matching method can only identify compounds within spectral database. In response, we present a deep-learning-based approach (DeepEI) for structure elucidation of unknown compound with its EI-MS spectrum. DeepEI employs deep neural networks to predict molecular fingerprint from EI-MS spectrum, and searches molecular structure database with the predicted fingerprints. In addition, a convolutional neural network was also trained to filter the structures in database and improve the identification performance. Our method shows improvement on the competing method NEIMS in identification accuracy on both NIST test dataset and MassBank dataset. Furthermore, DeepEI (spectrum to fingerprint) and NEIMS (fingerprint to spectrum) can be combined to improve identification accuracy.

## Introduction

Mass spectrometry is a widely used technique for identification compounds in biological systems [1–3]. GC-MS is particularly suitable for the detection of thermal-stable volatile compounds. Electron ionization (EI) is the most popular hard ionization method of GC-MS, which can generate extensive fragments for structure elucidation. Matching spectrum with database has been the routine method. The similarities are calculated between the spectrum of unknown and the spectra of standards in the database, and the structure of standard with the highest similarity is regarded as the structure of the unknown. Though widely used, it suffers from a coverage problem: if the compound is not in the database, it can never be identified correctly [4].

In-silicon fragmentation is an alternative approach to solve this problem. It calculates theoretical spectra of compounds from their chemical structures. Rule-based methods, quantum chemical calculation and machine learning are commonly strategies to calculate the theoretical spectra. Rule-based methods usually need experts to summarize the cleavage rules from large number of experimental spectra, which needs professional knowledge and efforts. Weissenberg and Dagan analyzed 190,825 entries in NIST 2005 database and found 68 ESI cleavage rules and their EI analogues [5]. Quantum chemical calculation methods use an equilibration and sampling strategy randomly selecting snapshots, then apply cascading production runs to generate fragments [6,7]. This approach has been demonstrated by a few representative spectra, but it needs high CPU computing power and running time. CFM-ID [8,9] is a machine learning method to predict ESI-MS/MS spectra, which applied a probabilistic generative model for evaluating the break tendency of chemical bonds, and trained the parameters of this model via machine learning approach. It can also be used for EI-MS, called CFM-EI [10], after some modification such as included odd electron fragment ions and isotope peaks. Recently, Wei et al proposed a neural network-based method, called NEIMS, which directly predicts EI-MS spectra from molecular fingerprints [4]. Comparing with CFM-EI, it not only achieves better predicting accuracy, but also runs much faster.

Apart from spectra prediction, there are also other strategies for compound identification. In order to rank the candidate structures based on the obtained spectra, one can predict the molecular fingerprints from the spectra. By comparing the predicted fingerprints against the structural database, the correct compound should be ranked at the top of the candidate list. This strategy was initially used for compound identification with ESI-MS/MS, called FingerID [11]. Subsequently, it was combined with multi-kernel learning and fragmentation tree for better performance [12,13]. Recently, MetExpert was developed to predict molecular fingerprints from EI-MS spectra, combined with retention index prediction and compound-likeness evaluation [14]. It achieved better accuracy than CFM-EI. However, only 188-bits MACCS (Molecular ACCess System) fingerprint [15] was used in MetExpert, which leads to the low discrimination degree of similar structures. Moreover, PLS-DA (partial least squares-discriminant analysis) model used by MetExpert is a linear classifier. In PLS-DA method, high-dimensions features are projected into a low-dimensional space to find a plane to differentiate different classes. Therefore, PLS-DA is not be optimal, if the fingerprint is not linear separable with the spectra.

In this study, a deep-learning-based approach (DeepEI) is proposed to retrieve the structure of the unknown compound from its EI-MS spectrum and the molecular structure database directly. Deep neural networks are used to predict multiple molecular fingerprints from the EI-MS spectrum (FP model). Then, the structure of the unknown compound can be retrieved by searching molecular structure database with the predicted fingerprints. In order to improve the identification performance, a convolutional neural network-based retention index model was trained to filter the structures in database. The overall workflow of DeepEI is described in Figure 1. DeepEI was tested by a subset of small molecules in NIST 2017 and an independent dataset from MassBank. The source code of DeepEI is available at https://github.com/hcji/DeepEI.

**Figure 1.**
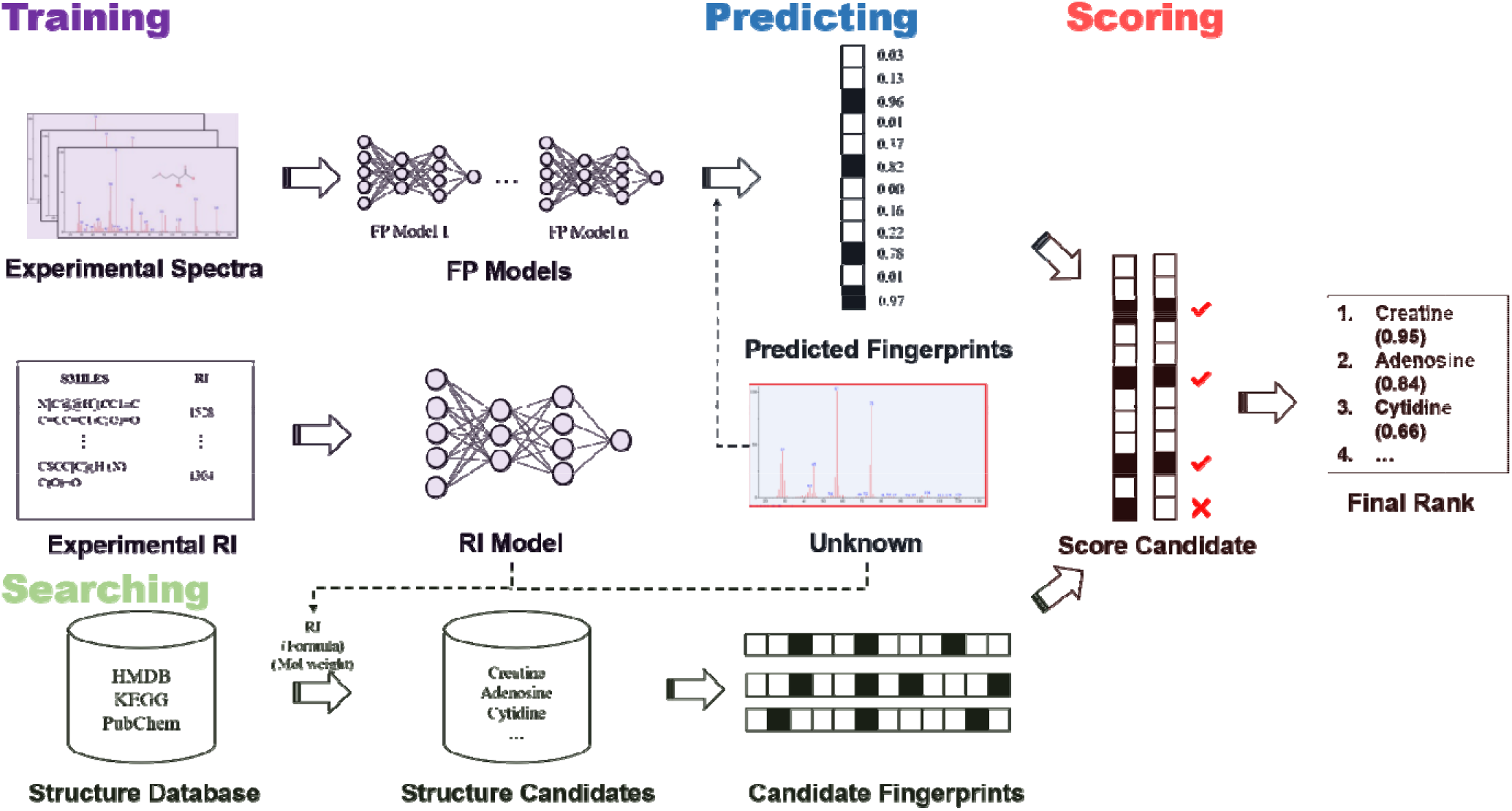
Workflow of the DeepMASS method. Spectra and retention indices in the database are used for training the RI model and fingerprint models, respectively. For spectrum and RI pair of an unknown, structure candidates can be retrieved with the RI. If the molecular weight is known, the candidate list can be shortened. The fingerprints of the unknown can be predicted by the FP model and the spectrum, and the fingerprints of the candidates can be calculated. By comparing with the predicted fingerprints and the calculated candidate fingerprints by a score function, the ranked candidate list can be obtained.

## Methods

### Datasets and Preparation

The spectra used in this study are from two databases: NIST 2017 and MassBank. Spectra were obtained from NIST 2017 main library. The duplications, the compounds with molecular weight over 2000 Da or uncommon elements (elements except C, H, O, N, P, S, F, Cl, Br, Si) were excluded. Finally, 195,285 spectra were remained. A subset of 10,411 spectra were split for independent test for evaluating the performance of the compound identification, and it is the same as the independent test set in NEIMS for fairly comparison in the subsequent procedure.

Spectra of MassBank were downloaded from the homepage of the MS-DIAL software [16,17]. The msp file were parser by MetGem package [18]. SMILES of a few compounds cannot be parsed by *rdkit*, and they were ignored. Finally, 13,083 spectra were obtained, and all of them were used as external test dataset.

### Molecular Fingerprints Prediction (FP model)

The abundant fragments in EI-MS spectrum bring the possibility to predict the molecular fingerprint from the spectrum directly. We trained full connected neural network model for each bit of fingerprint, which represented the substructure information of the unknown compound. In this work, six kinds of substructure-related molecular fingerprints were used and listed in Table 1 [15,19–23]. There are 8034 fingerprint bits when combining all the six kinds of molecular fingerprints. However, most of the fingerprint bits are class-unbalanced. We only kept the fingerprint bit with positive percentage between 10 % and 90 %, and 636 bits of fingerprints were obtained finally.

**Table 1.** Types of fingerprints used by DeepEI.

| **Fingerprint type** | **Bits** | **Reference** |
| --- | --- | --- |
| CDK standard fingerprint | 1024 | [19] |
| PubChem fingerprint | 881 | [20] |
| Klekota-Roth fingerprint | 4860 | [21] |
| MACCS keys | 166 | [15] |
| Estate fingerprint | 79 | [22] |
| Circular fingerprint | 1024 | [23] |

Since the remaining spectra in NIST 2017 are integral mass-to-charge value with molecular weights less than 2000 Da. As shown in Figure 2, all the spectra were converted to vectors with size=2000 as the inputs of the neutral networks. Each model for molecular fingerprint prediction contains three fully-connected layers (units=2000, 1000 and 500 respectively) with ReLU activation function. The output layer is a fully-connected layer (units=2) with SoftMax activation function. Categorical cross-entropy was used as the loss function, and Adam optimization algorithm was used as the optimizer. 8 epochs were performed on the training dataset.

**Figure 2.**
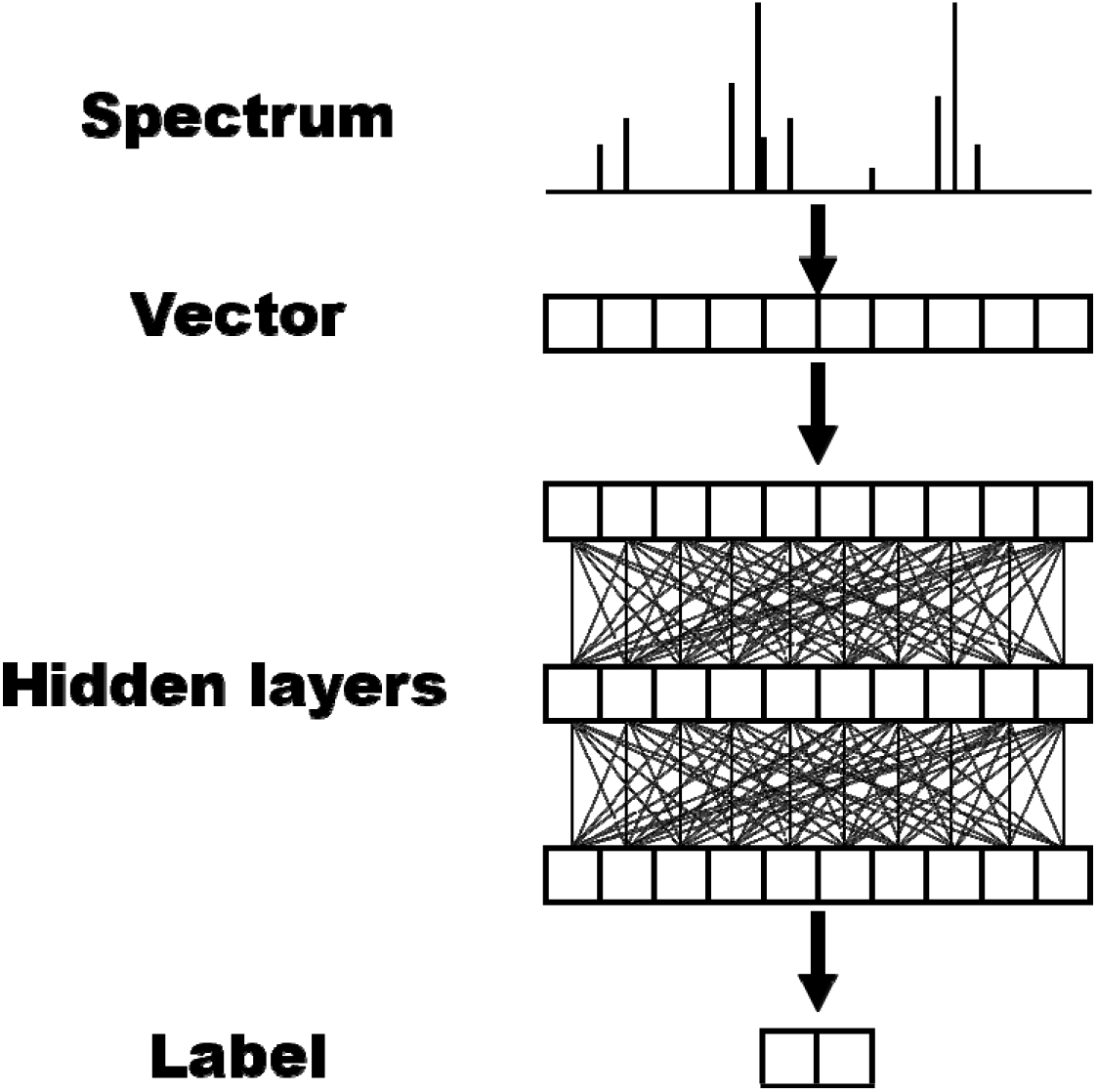
The neural network structure of FP model in DeepEI.

### Compound Identification

After training the FP model models, the molecular fingerprints can be predicted for given spectrum of unknown. In order to annotate its structure, the candidate structures need to be retrieved from the compound database and filtered by the retention indices and/or the molecular weight. In DeepEI, a convolutional neural network-based retention index predicting model (RI model) was provided to predict the retention index of compounds in the database, filter structures in the database and obtain the structure candidates. Five different deep-learning-based RI models were trained and compared, and multi-channel CNN have the best performance. The detailed results can be found in the “*Retention Index Prediction*” section of support information. Next, the fingerprints of the candidate structures are computed and compared with the predicted fingerprints of the unknown using the Jaccard function. Finally, the ranked candidate list can be obtained by sorting the Jaccard similarity scores in descending order, which defined as follow:

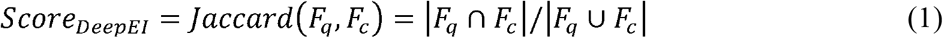

where *F_q_* represents the query fingerprints predicted by DeepEI, and *F_l_* represents the fingerprints of the candidates in molecular structure library.

## Results

### Evaluation of FP Model

We trained totally 636 FP models and each model predicted a fingerprint bit related to the substructure of the compounds. The compounds were split to train, valid and test set. We also compared the performance of the FP model with the partial least squares-discriminant analysis (PLS-DA), logistic regression (LR) and gradient boosting (XGBoost). In the PLS-DA method, the number of components were optimized for the best performance on the validation set. The results of test set are shown in Figure 3. From the results, the median predicting accuracy of FP models is 90.3 %, which indicates the substructure information of the compounds can be well predicted by these models. The median F1 score of the FP models is also over 79.2 %, which indicates the FP models are reliable in the prediction of fingerprint bits. PLS-DA, LR and XGBoost also achieve average accuracy of 82.3 %, 83.2 % and 86.4 %, respectively. However, their average F1 scores are much lower than DeepEI, which are 37.1 %, 46.3 % and 58.8 %, respectively. It means the predicting results of DeepEI are more meaningful.

**Figure 3.**
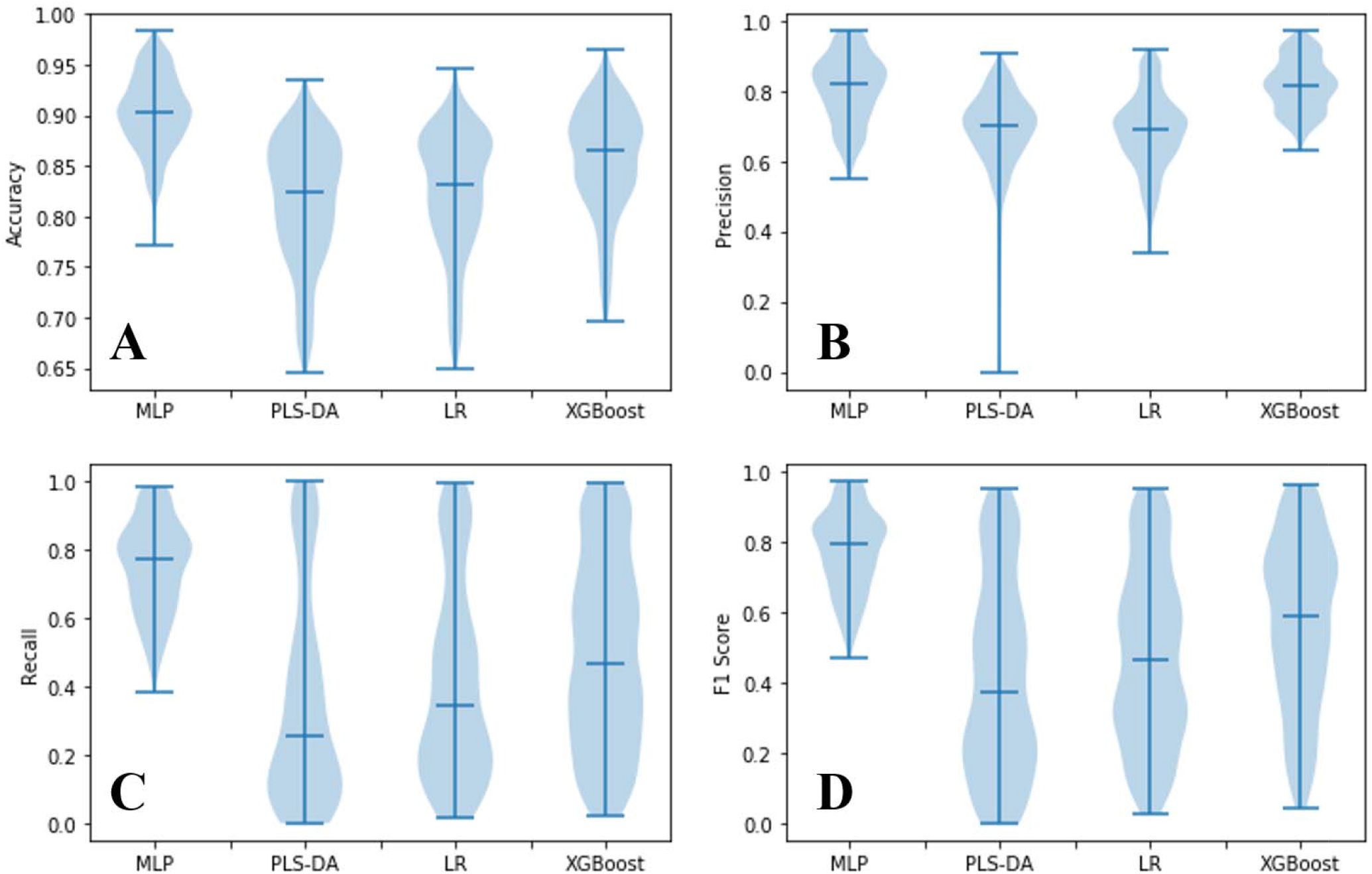
Comparison of fingerprint prediction of accuracy (A), precision (B), recall (C) and F1 score (D) of MLP, PLS-DA, LR and XGBoost method.

### Compound Identification Performance

We compared the compound identification performance of DeepEI and NEIMS. NEIMS is a recently published method, which predicts EI mass spectra for enhance compound identification with neural network [4]. The model and the source code of NEIMS were download from https://github.com/brain-research/deep-molecular-massspec. The evaluation was performed with the NIST subset and the MassBank dataset. These compounds were never used for training, parameter tuning or model selection. Each compound with the experimental EI-MS spectrum was treated as the “unknown” to evaluate the identification performance. Candidate structures were retrieved from all compounds in NIST database. The 5 Da molecular weight filter was used, which means the molecular weights difference between the candidate structures and the “unknown” compound should be less than 5 Da.

As described in the reference, the augmented library by NEIMS was used for identification. The spectrum of the “unknown” was predicted and inserted to the candidate list. If the “unknown” compound was already in the NIST database, we replaced the original experimental spectrum with the predicted spectrum. Then we used the candidate list for spectra matching. The weighted cosine similarity was used for ranking the candidates:

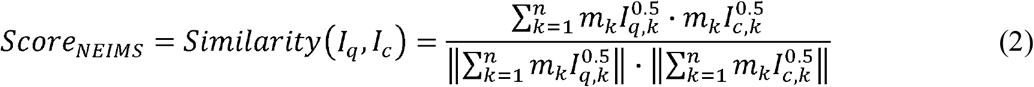

where *I_q_* and *I_c_* are vectors of m/z intensities representing the query spectrum and the candidate spectrum, respectively. *m_k_* and *I_k_* are the m/z value and intensity found at m/z. n is the largest m/z index.

The results are listed in Table 2. For the NIST subset, the recall@1 and recall@5 of DeepEI are 36.8 % and 66.2 %, respectively. While the recall@5 of DeepEI is similar to NEIMS, the recall@1 is much higher. For the MassBank dataset, the recall@1 and recall@5 are 43.0 % and 57.5 %. The results are both much higher than NEIMS. The detailed identification results are available as support information. Overall, DeepEI is very effective for identifying the correct structure of the unknown.

**Table 2.** Comparison between DeepEI and NEIMS with NIST and MassBank dataset.

| **Test set** | **Rank** | **DeepEI** | **NEIMS** | **DeepEI+NEIMS** |
| --- | --- | --- | --- | --- |
| NIST 2017 | Top 1 | 36.8 % | 27.2 % | 37.1 % |
|  | Top 3 | 58.2 % | 54.6 % | 70.1 % |
|  | Top 5 | 66.2 % | 67.8 % | 82.5 % |
| MassBank | Top 1 | 43.0 % | 22.6 % | 32.2 % |
|  | Top 3 | 53.3 % | 41.4 % | 56.4 % |
|  | Top 5 | 57.5 % | 50.6 % | 65.1 % |

### Combination with NEIMS

DeepEI is a spectrum to fingerprint method and NEIMS is fingerprint to spectrum method, they were supposed to be complementary for each other and may achieve a better identification performance if used together. Therefore, a consensus score was proposed by weighted averaging the scores of NEIMS and DeepEI. We also used the NIST subset and the MassBank dataset to compare the identification performance using the consensus score, which can be defined as:

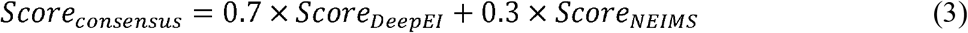

As shown in Table 2, The results of DeepEI + NEIMS have a significant improvement comparing with the results of NEIMS alone. Also, DeepEI + NEIMS have much better recall@1, recall@3 and recall@5 on NIST 2017 dataset, and better recall@3 and recall@5 on MassBank dataset.

## Conclusion

In this work, we use deep neutral networks to predict retention index and molecular fingerprint to assist compound identification with EI-MS. By comparing three FCNN networks and two CNN networks with different kinds of inputs, the multi-channel CNN achieved the best performance in predicting retention index of all kinds of chromatographic columns. We also compared PLS-DA, LR, XGBoost and FP model for molecular fingerprint prediction. The results indicated FP model has much better performance. The compound identification performance of DeepEI are evaluated by test dataset of NIST2017 and MassBank dataset. Results show that DeepEI is effective for identifying the correct structure of the unknown. Compared with NEIMS on NIST test dataset, DeepEI has a similar performance on recall@5, but significant better performance on recall@1 and recall@3. Compared with NEIMS on MassBank dataset, DeepEI has significant better performance on all the recall@1, recall@3 and recall@5. Meanwhile, the combination of DeepEI and NEIMS can improve the compound identification accuracy by a significant margin because of their complementarity.

## Supporting information

support information

## Acknowledgements

This work is financially supported by the National Natural Science Foundation of China (Grant Numbers. 21873116 and 21675174)

## TOC

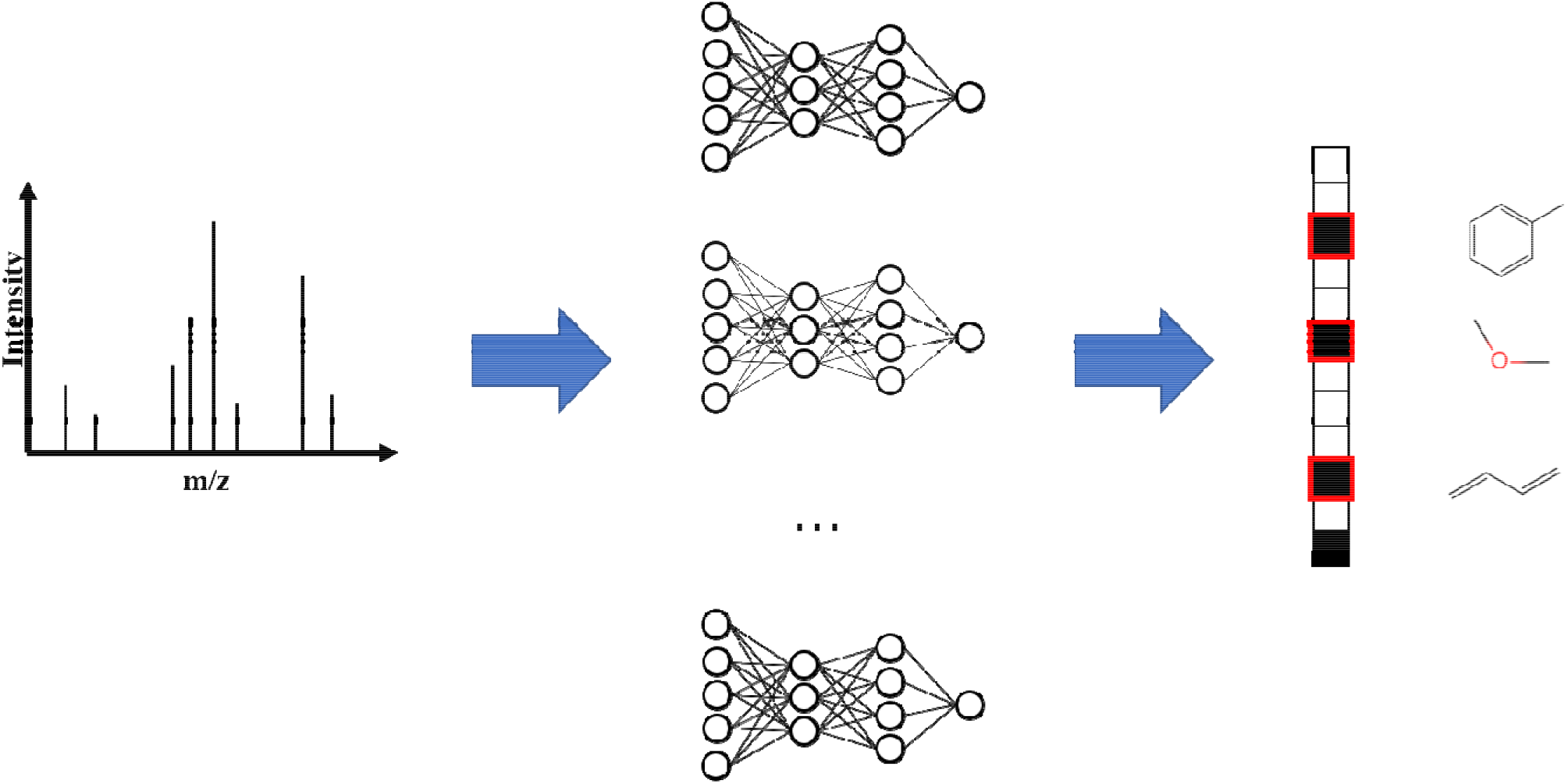

