## Supplementary material for "Predicting Molecular Fingerprint from Electron−Ionization Mass Spectrum with Deep Neural Networks": support information

**Retention Index Prediction**

Machine learning based prediction of GC retention index were previous reported. The input features include chemical descriptor [1], molecular fingerprints [2] and SMILES strings [3]. In this work, we compared five different neural networks and evaluated the performance of the models. The details of each neutral networks are summarized in Figure S1. Two multilayer perceptron (MLP) used Morgan fingerprints [4], chemical descriptors. The Morgan fingerprints were calculated by RDKit package (https://rdkit.org), and all the chemical descriptors were calculated by RCDK package [5]. In each MLP, five fully-connected layers were followed by the input layer, and each layer used the ReLU function as the activation function. The third MLP concatenates the fingerprints and the chemical descriptors first, then follows a series of fully-connected layers. Two convolutional neutral networks (CNN) used the one-hot encoded SMILES strings as the input features [6]. The first network is called single-channel CNN. In this network, five one-dimension convolutional layers, with stride 1, kernel 4, were followed by the input layer, and followed by a fully-connected layer. The second network is called multi-channel CNN. In this network, three one-dimension convolutional layers with different kernel sizes were followed by the input layer respectively. Each convolutional layer was followed by another convolutional layer. Then, the outputs of the convolutional layers were concatenated and followed by a fully-connected layer. All the layers in the CNNs used a rectified linear unit (ReLU) function as the activation function. The other hyper-parameters were optimized by the valid dataset.

The goal was to build a regression model minimizing the error between the prediction retention index and the experimental retention index. Thus, the linear activation function was used in the output layer and the mean square error was used as the loss function. The Adam optimization algorithm was used for training the deep neural networks. 20 epochs were performed on the training dataset. In order to avoid overfitting, we also used an early stopping method with the monitor of the loss of valid dataset.

**Evaluation of the RI Model.** In the NIST 2017 database, not all of the compounds have the retention index values. Among the included compounds of this work, 50094 compounds have RI values of the semi-standard nonpolar column (SemiStdNP); 9609 and 5353 compounds have RI values for the standard nonpolar column (StdNP) and the standard polar column (StdPolar), respectively. When training the RI models, only the compounds with RI values were used. They were split into train, valid and test sets, and the split ratio is 8:1:1. The test set was used for evaluating the performance of the RI models. Figure S2 shows the performances of different models for different kinds of chromatographic columns. From the results, MLP model with molecular descriptors has pretty good performance in predicting RI of SemiStdNP column, but the performance on StdNP column and StdPolar are unsatisfactory. MLP model with fingerprints work the worst in the compared methods. However, if combined the fingerprints with the molecular descriptors, the performance is significant improved. CNN models can take one-hot encoded SMILES strings as input directly, which avoid the computation of molecular descriptors. Comparing with single-channel CNN, multi-channel CNN shows significant better performance on StdNP and StdPolar. In summary, multi-channel CNN is the best model in the compared models for RI prediction.

**RI Filter for Identification.** We applied the multi-channel CNN RI model to improve the metabolite identification performance. Since not all compounds in NIST 2017 have RI, only those with RI of SemiStdNP column was evaluated. The results are listed in Table S1. When no filter was used, the recall@1 and recall@5 were 25.7 % and 46.1%. When RI filter was used, the values promote into 28.2 % and 51.2 %, respectively. Although with mass filter, the recall@1 and recall@5 are higher, it is difficult to obtain the exact masses of metabolites from GC-EI-MS. Therefore, the RI model and filter were effective and practical for metabolite identification.

Table S1. Improve the compound identification performance of DeepEI with RI filter.

| **Rank** | **Without filter** | **RI filter** | **Mass filter** |
| --- | --- | --- | --- |
| Top 1 | 25.7 % | 28.2 % | 49.7 % |
| Top 3 | 39.3 % | 43.2 % | 61.4 % |
| Top 5 | 46.1 % | 51.2 % | 70.0 % |

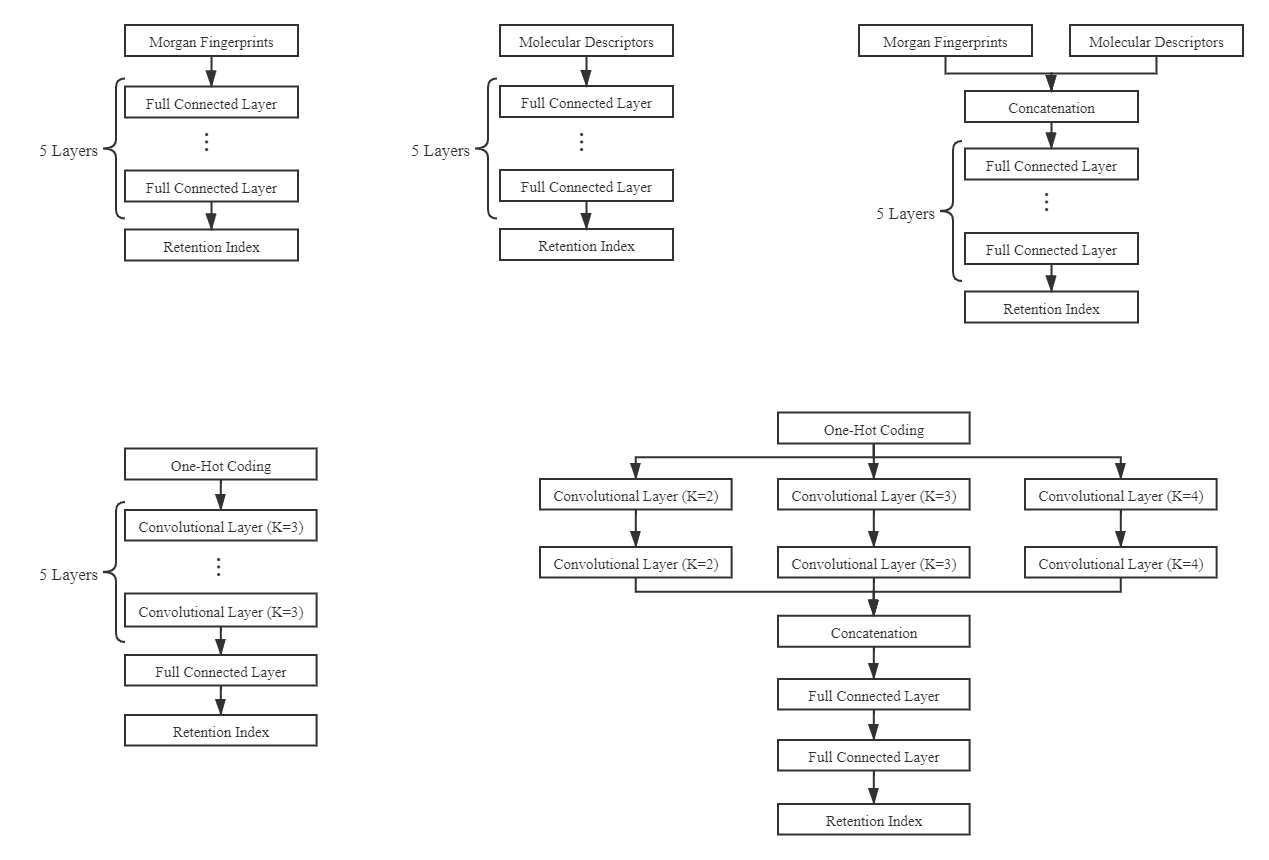
Figure S1. Structure of the neural networks: MLP-Fingerprints, MLP-Descriptors, MLP-Combination, single-channel CNN, multi-channel CNN.

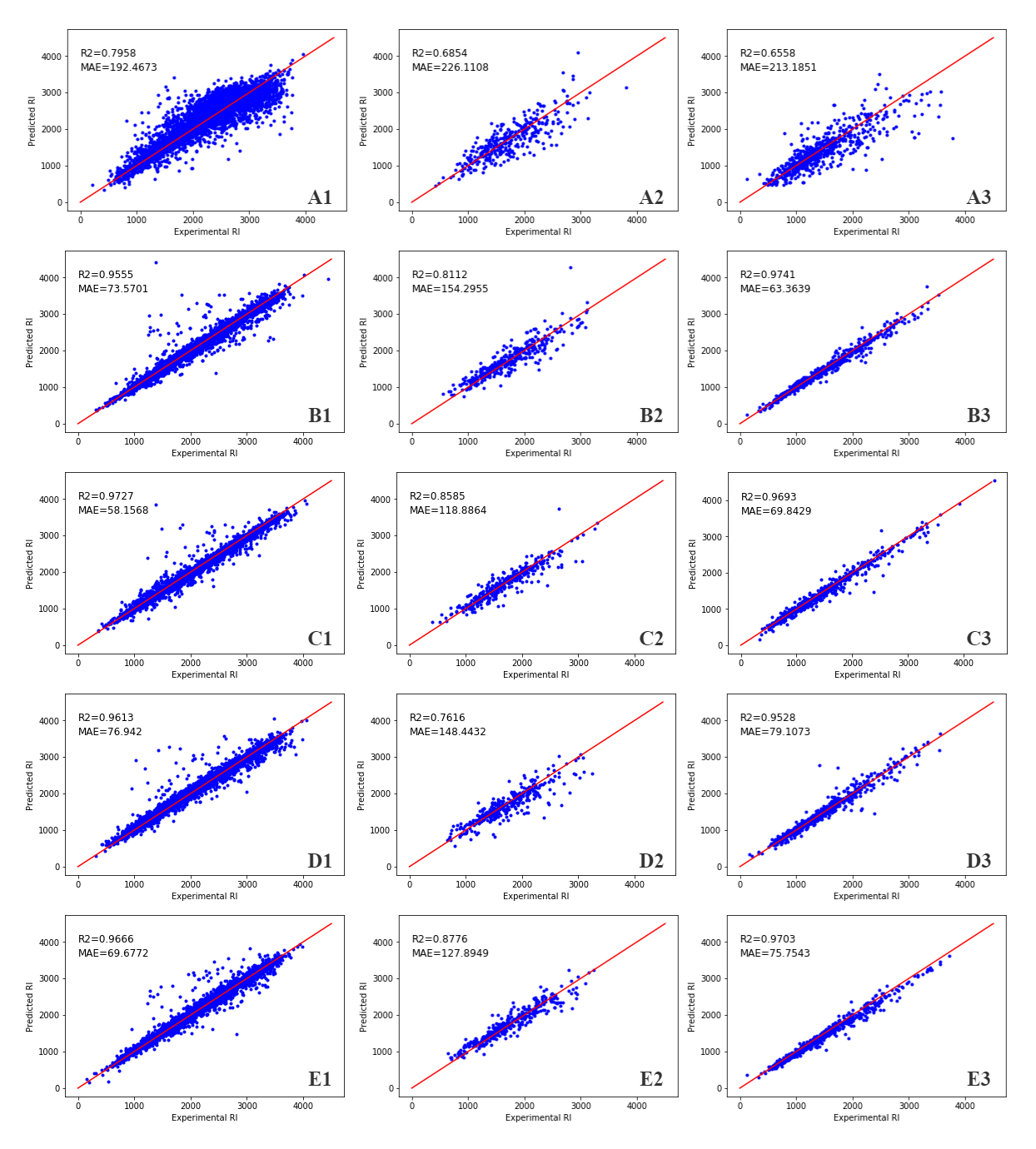
Figure S2. In this figure, the letter A-E is the models corresponding to MLP-Fingerprints, MLP-Descriptors, MLP-Combination, single-channel CNN, multi-channel CNN, while the number 1-3 is the chromatographic columns corresponding to semi-standard nonpolar column (SemiStdNP), the standard polar column (StdPolar) and standard nonpolar column (StdNP).

1. College of Chemistry and Chemical Engineering, Central South University, Changsha 410083, PR China. Correspondence and requests for materials should be addressed to H.L. and Z.Z.
